# Analysis on epidemiological characteristics of 4 species natural-focal diseases in Shandong Province, China in 2009-2017: a descriptive analysis

**DOI:** 10.1101/564971

**Authors:** Rui Chen, Zengqiang Kou, Liuchen Xu, Jie Cao, Ziwei Liu, Xiaojing Wen, Zhiyu Wang, Hongling Wen

## Abstract

**Background:** Natural-focal diseases have always been a kind of serious disease that endangers human health. It threatens about 100 million people in Shandong Province, and causes illness in thousands of people each year. However, information on the epidemiological characteristics of natural-focal diseases in Shandong Province has been limited. The purpose of the study is to describe and analyze the epidemiological characteristics of natural-focal diseases in Shandong Province, 2009-2017.

**Methods:** We describe the incidence and distribution of 4 species natural-focal diseases in Shandong Province using surveillance data from 2009-2017.

**Results:** From 2009-2017, 11123 cases of 4 species natural-focal diseases including 257 deaths were reported in Shandong Province, China. The 4 species natural-focal diseases were severe fever with thrombocytopenia syndrome (SFTS), human granulocytic anaplasmosis (HGA), typhus and scrub typhus respectively. The high-risk groups of the 4 species diseases all were farmers and the elderly. The incidence rate of scrub typhus was significantly higher in females, however, this difference was not seen in the other 3 diseases. The 4 species diseases were mainly clustered in middle-southern part of Shandong (Mount Yimeng) and Shandong Peninsula (Laoshan Mountain). The annual incidence of SFTS and scrub typhus had increased in the mass, typhus had been relatively stable, and HGA had declined. However, the popular range of SFTS had been expanding, HGA had been shrinking, and typhus and scrub typhus were unchanged. The epidemic period of SFTS and HGA was from May to October, typhus was from October to November, scrub typhus was from September to November in Shandong Province. The fatality rates of SFTS, typhus, scrub typhus, HGA were 9.19%, 0%, 0.01%, 2.24%, respectively.

**Conclusions:** Our study described and analyzed the prevalence of natural-focal diseases in Shandong, and confirmed that age was closely related to the SFTS fatality rate. This study may be applicable to an improved understanding of the prevalence of natural-focal diseases in Shandong Province in recent years and the better development of the accurate prevention and control strategies for natural-focal diseases in Shandong Province.

**Author Summary:** Natural-focal diseases are a serious public health problem in Shandong Province, China. It threatens about 100 million people in Shandong Province, and causes illness in thousands of people each year. This study used the monitoring data from Shandong Province for 2009-2017 to describe and analyze the epidemiological characteristics of the 4 species natural-focal diseases. The results showed that the risk population of natural-focal diseases in Shandong Province was farmers and the elderly, the epidemic season was mainly in summer and autumn, the middle-southern part of Shandong Province and Shandong Peninsula were more seriously affected compared with other regions. In addition, the epidemic area of SFTS was expanded, with a fatality rate of 9.19%. These findings indicated that public awareness of natural-focal diseases should be raised in the epidemic focus, especially for farmers, and further efforts should be strengthen specially in high-risk areas and during the epidemic season.

## Introduction

Natural-focal diseases are a large group of diseases. At present, there are more than 180 kinds of natural-focal diseases in the world, including viral diseases, bacterial diseases, rickettsiosis, chlamydia, spirochetes, fungal diseases, protozoa diseases and other parasitic diseases. However, in recent years, a variety of old natural-focal diseases have revived in the world, such as Ebola virus disease (EVD), Zika virus disease, dengue fever(DF), etc. There were 28,646 reported cases (suspected, probable, and confirmed) for only EVD in 2014-2016 with 11,323 deaths reported in West Africa as of March 30, 2016[1]. In addition, new pathogens of natural-focal diseases have constantly been found in China, such as severe fever with thrombocytopenia syndrome phlebovirus(SFTSV), Candidatus Rickettsia tarasevichiae, Rickettsia sibirica Subspecies sibirica BJ-90, etc[2-4]. In the past 55 years, about 8,350,754 cases of natural-focal diseases involving 24 types of natural-focal diseases were reported in Chinese journals only[5]. Natural-focal diseases constitute a serious threat to the public health. As a large agricultural and population province in China, Shandong Province has more than a 40 million agricultural population. Most of the young people study and work in cities in addition to the Spring Festival. Most older people and children live in rural areas all year round. The climate in Shandong Province is warm with four distinct seasons and the terrain is diverse, which provides favorable conditions for the occurrence of many natural-focal diseases. In the capital city of Jinan alone, there were 1248 reported cases for 9 different natural-focal diseases in 2004-2013[6]. However, limited information concerning the epidemiologic characteristics of natural-focal diseases in Shandong Province is available, especially the systematic description of this a large group of diseases. Therefore, it is necessary to systematically describe the epidemic characteristics of the natural-focal diseases in Shandong Province in recent years.

In order to make the study of the epidemic situation of natural-focal diseases more representative, we chose 4 species natural-focal diseases with low to high incidence levels to represent the overall incidence of natural-focal diseases in Shandong Province. At the same time, these 4 species natural-focal diseases also represent 3 different kinds of insect borne diseases. The incidence rate levels from low to high are HGA, typhus, SFTS, and scrub typhus, respectively. The pathogens of SFTS and HGA are SFTSV and Anaplasma phagocytophilum (AP), respectively, and the main vectors are ticks[7]. The Typhus, including epidemic typhus and endemic typhus, is caused by Rickettsia typhus. The typhus in Shandong is mainly endemic typhus [8]. Endemic typhus is caused by Ricettsia mooseri and is mainly transmitted by Xenopsyllae cheopis, rodents are their natural hosts. Scrub typhus is caused by the intracellular pathogen Orientia tsutsugamushi and is transmitted by chigger mites[9]. We describe the magnitude and distribution of these diseases in Shandong Province based on the notifiable reporting date, focusing on three-dimensional distribution from 2009 to 2017 and characteristic of SFTS fatality. SFTS was not monitored until 2010 in China, hence there was no monitoring data for SFTS in 2009. There were few cases reported for HGA in Shandong Province in recent years, and HGA was not monitored in 2017.

## Methods

### Study site

Shandong Province is located on the eastern coast of China (longitude 114°19’ E to 122°43’ E, latitude 34°22’ N to 38°23’ N), with an area of 157,000 km^2^, of which the vegetation coverage is about 77.54%, the planted land area is about 53.82%, the mountain area is about 14.59%, and the hilly area is about 15.39%. It is mountainous in the middle-southern part of Shandong (Mount Yimeng) and Shandong Peninsula (Laoshan Mountain) but mostly flat and hilly in the periphery. It is in the lower reaches of the Yellow River, and it extends out to the Pacific Ocean in the form of the Shandong Peninsula, with a coastline of 3,121 km (Fig 1). Shandong Province is the second most populous province in China with a population of about 100 million, of which about 40% is agricultural. Shandong has a monsoon climate of medium latitudes (average annual temperature is 13.6–14.3°C; average annual precipitation is 543–845 mm).

**Fig 1.**
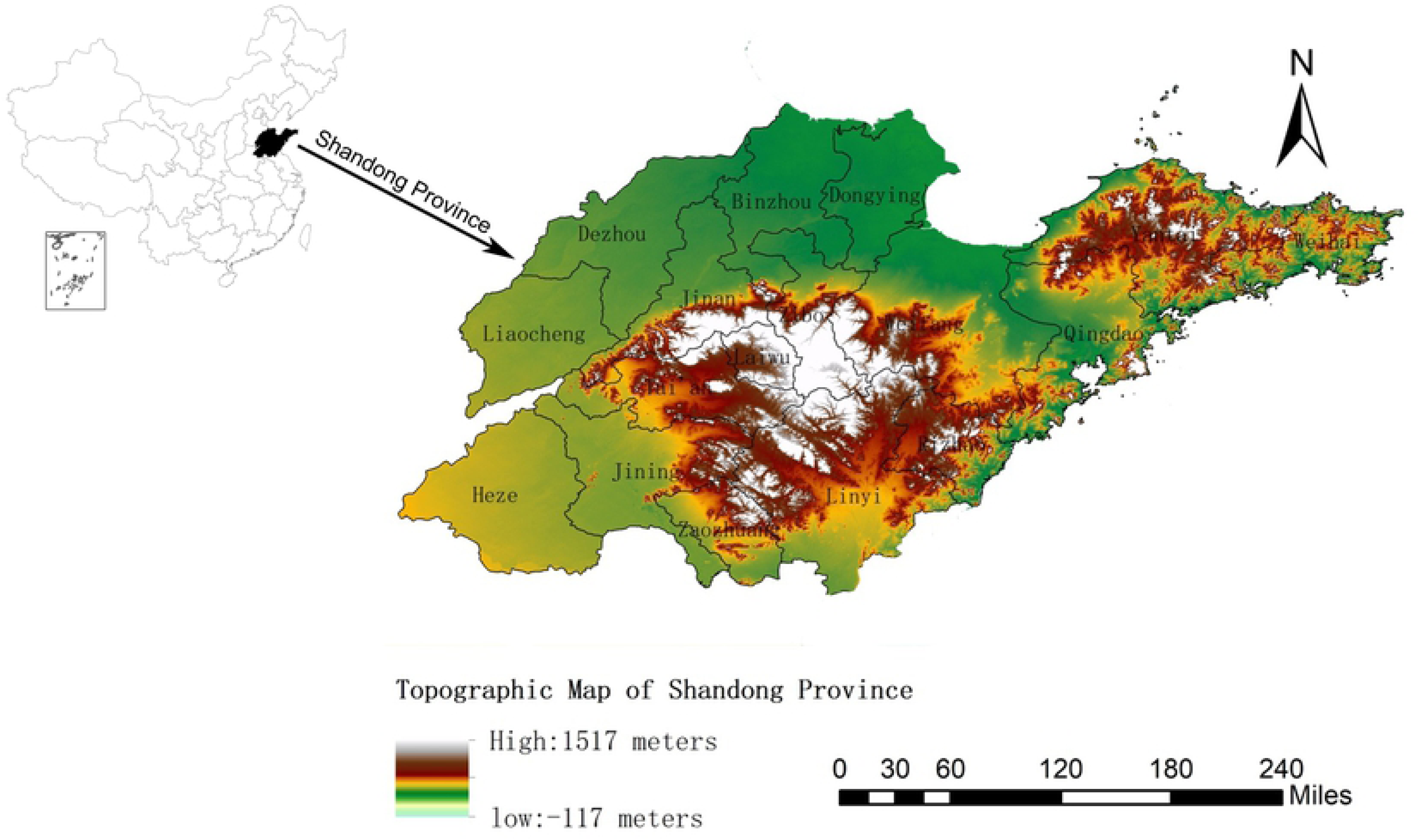
Topographic map of Shandong Province.

### Case definition

Since January 1, 2004, China has started to use the network direct reporting system of China Infectious Disease Reporting Information System. 4 species natural-focal diseases cases are diagnosed according to the diagnosis criteria issued by the health department of China (S2 Appendix).

### Data collection

Daily disease surveillance data on SFTS, typhus, scrub typhus and HGA in Shandong Province from 2009 to 2017 were obtained from the Shandong Disease Reporting Information System (SDRIS). Information of 4 species of natural-focal diseases cases included gender, age, occupation, residential address, date of illness onset and outcome of illness. Export the infectious disease report card according to the date of onset. Population data for 2009-2017 of Shandong Province came from Shandong Province Bureau of Statistics.

### Data analysis

The age distribution, gender distribution, occupation distribution, seasonal distribution and regional distribution of the cases were summarized using Excel 2010, and IBM SPSS Statistics 24.0 (online) was used to perform the statistical analysis. The different rates were analyzed using the *χ*^*2*^ test. Trend test was analyzed using the Linear-by-linear Association. All tests were 2-tailed and statistical significance was set at *P* < 0.05. The base diagrams in Figure 1 and Figure 6 were from the Resource and Environment Data Cloud Platform. The software ArcGis10.2 was used to plot the topographic map of Shandong Province and the geographical distribution of cases (Fig 1 and Fig 6).

### Ethical approval

It was determined by the National Health and Family Planning Commission, China, that the collection of data from natural-focal diseases cases was part of a continuing public health surveillance of a notifiable infectious and was exempt from institutional review board assessment.

## Results

### SFTS

From 2010 to 2017, a total of 2731 confirmed cases of SFTS including 251 deaths in Shandong Province were reported to SDRIS.

#### Time distribution

During 2010-2017, the number of SFTS had been on the rise in the mass except that in 2012 and 2017 the number declined slightly, with the highest recorded in 2016 (629 cases) (Fig 2B). An average of 3.47 cases per one million residents were reported each year in Shandong Province during 2010-2017, with the highest recorded in 2016 (6.32 cases/1,000,000) and the lowest in 2010 (0.59 cases/1,000,000) (Fig 2A). The incidence of SFTS showed obvious seasonal characteristics. During 2010–2017, 94.03% (2568/2731) of SFTS cases were reported from May to October, the peaks of reported SFTS cases from 2010 to 2017 occurred in the summer season, July, July, July, June, June, May, June, August, respectively (Fig 3A; Table 1).

**Table 1.** Demographic and epidemiological characteristics of 4 species of natural-focal diseases in Shandong province, 2009-2017.

|  | SFTS<br>(2010-2017) |  | HGA<br>(2009-2016) |  | Typhus<br>(2009-2017) |  | Scrub typhus<br>(2009-2017) |  |
| --- | --- | --- | --- | --- | --- | --- | --- | --- |
|  | No. of<br>cases | Proportio<br>n (%) | No. of<br>cases | Proportio<br>n (%) | No. of<br>cases | Proportio<br>n (%) | No. of<br>cases | Proportio<br>n (%) |
| Age group (years) |  |  |  |  |  |  |  |  |
| 0- | 3 | 0.11 | 0 | 0.00 | 19 | 2.09 | 116 | 1.60 |
| 5- | 0 | 0.00 | 1 | 0.45 | 14 | 1.54 | 59 | 0.81 |
| 10- | 6 | 0.22 | 0 | 0.00 | 23 | 2.53 | 58 | 0.80 |
| 15- | 9 | 0.33 | 0 | 0.00 | 11 | 1.21 | 29 | 0.40 |
| 20- | 9 | 0.33 | 2 | 0.90 | 19 | 2.09 | 50 | 0.69 |
| 25- | 36 | 1.32 | 1 | 0.45 | 20 | 2.20 | 141 | 1.94 |
| 30- | 34 | 1.24 | 6 | 2.69 | 15 | 1.65 | 167 | 2.30 |
| 35- | 43 | 1.57 | 7 | 3.14 | 46 | 5.06 | 213 | 2.93 |
| 40- | 104 | 3.81 | 13 | 5.83 | 68 | 7.48 | 429 | 5.91 |
| 45- | 193 | 7.07 | 21 | 9.42 | 85 | 9.35 | 671 | 9.24 |
| 50- | 327 | 11.97 | 20 | 8.97 | 108 | 11.88 | 1012 | 13.94 |
| 55- | 383 | 14.02 | 30 | 13.45 | 122 | 13.42 | 1049 | 14.45 |
| 60- | 439 | 16.07 | 25 | 11.21 | 128 | 14.08 | 1053 | 14.50 |
| 65- | 420 | 15.38 | 33 | 14.80 | 86 | 9.46 | 864 | 11.90 |
| 70- | 336 | 12.30 | 30 | 13.45 | 69 | 7.59 | 618 | 8.51 |
| 75- | 225 | 8.24 | 24 | 10.76 | 45 | 4.95 | 410 | 5.65 |
| 80- | 128 | 4.69 | 9 | 4.04 | 25 | 2.75 | 238 | 3.28 |
| >85 | 36 | 1.32 | 1 | 0.45 | 6 | 0.66 | 83 | 1.14 |
| Total | 2731 | 100.00 | 223 | 100.00 | 909 | 100.00 | 7260 | 100.00 |
| Gender |  |  |  |  |  |  |  |  |
| Male | 1384 | 50.68 | 114 | 51.12 | 482 | 53.03 | 3148 | 43.36 |
| Female | 1347 | 49.32 | 109 | 48.88 | 427 | 46.97 | 4112 | 56.64 |
| Occupation |  |  |  |  |  |  |  |  |
| Child | 3 | 0.11 | 0 | 0.00 | 22 | 2.42 | 146 | 2.01 |
| Student | 14 | 0.51 | 1 | 0.45 | 40 | 4.40 | 106 | 1.46 |
| Laborer | 51 | 1.87 | 7 | 3.14 | 29 | 3.19 | 166 | 2.29 |
| Farmer | 2349 | 86.01 | 186 | 83.41 | 773 | 85.04 | 6420 | 88.43 |
| Retiree | 77 | 2.82 | 12 | 5.38 | 14 | 1.54 | 120 | 1.65 |
| Other | 237 | 8.68 | 17 | 7.62 | 31 | 3.41 | 302 | 4.16 |
| Total | 2731 | 100.00 | 223 | 100.00 | 909 | 100.00 | 7260 | 100.00 |
| Month |  |  |  |  |  |  |  |  |
| Jan | 8 | 0.29 | 0 | 0.00 | 17 | 1.87 | 9 | 0.12 |
| Feb | 8 | 0.29 | 2 | 0.90 | 7 | 0.77 | 3 | 0.04 |
| Mar | 13 | 0.48 | 0 | 0.00 | 11 | 1.21 | 6 | 0.08 |
| Apr | 73 | 2.67 | 4 | 1.79 | 16 | 1.76 | 12 | 0.17 |
| May | 472 | 17.28 | 24 | 10.76 | 68 | 7.48 | 47 | 0.65 |
| Jun | 549 | 20.10 | 40 | 17.94 | 34 | 3.74 | 34 | 0.47 |
| Jul | 532 | 19.48 | 42 | 18.83 | 35 | 3.85 | 29 | 0.40 |
| Aug | 538 | 19.70 | 53 | 23.77 | 31 | 3.41 | 48 | 0.66 |
| Sept | 273 | 10.00 | 32 | 14.35 | 44 | 4.84 | 370 | 5.10 |
| Oct | 204 | 7.47 | 20 | 8.97 | 517 | 56.88 | 5235 | 72.11 |
| Nov | 45 | 1.65 | 4 | 1.79 | 113 | 12.43 | 1442 | 19.86 |
| Dec | 16 | 0.59 | 2 | 0.90 | 16 | 1.76 | 25 | 0.34 |

**Fig 2.**
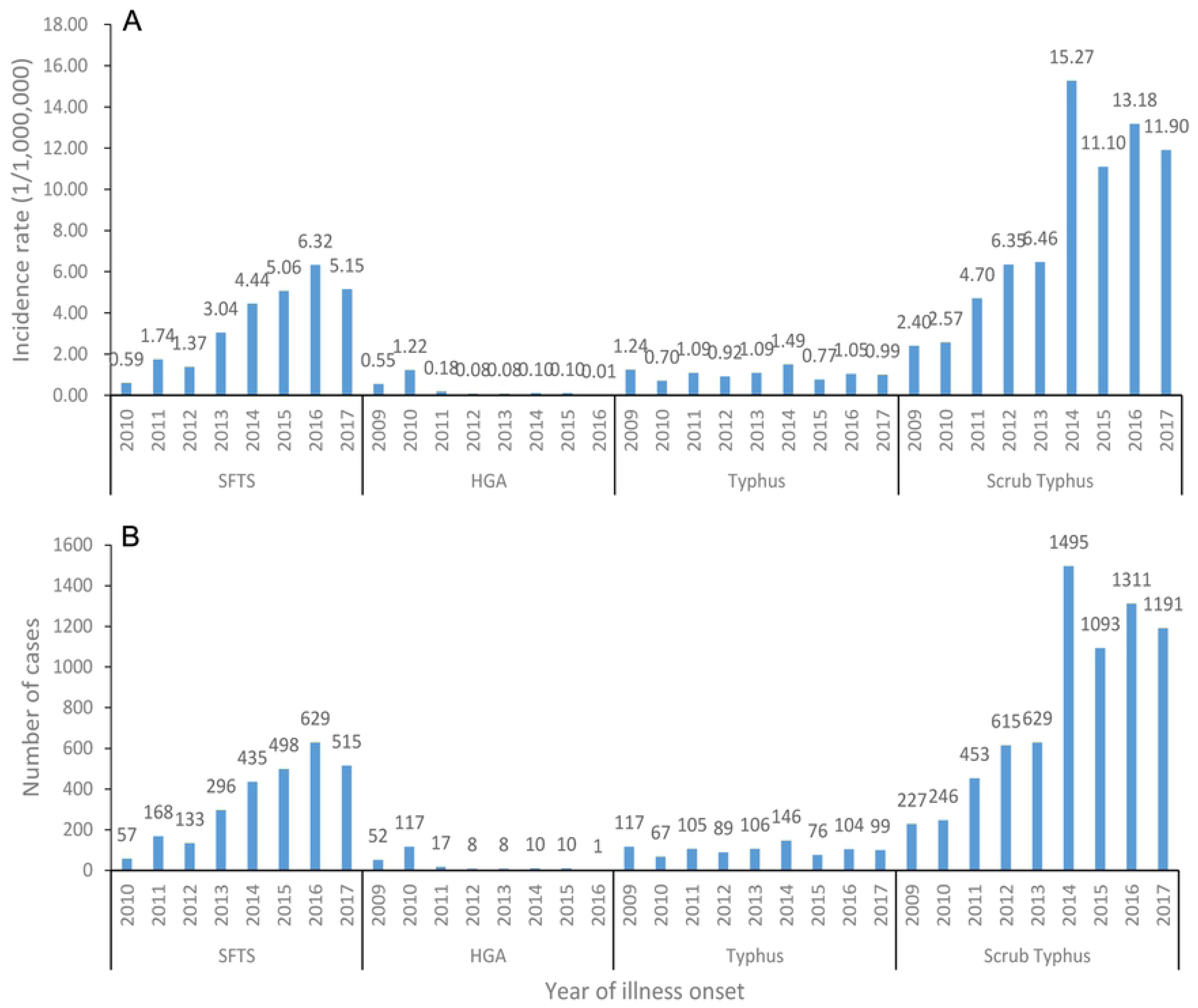
During 2009-2017, the incidence of 4 species natural-focal diseases in Shandong Province. (A): The morbidity of SFTS, HGA, typhus and scrub typhus per one million residents of Shandong Province at the end of each year. (B): The aggregated number of cases of SFTS, HGA, typhus and scrub typhus by year in Shandong Province.

**Fig 3.**
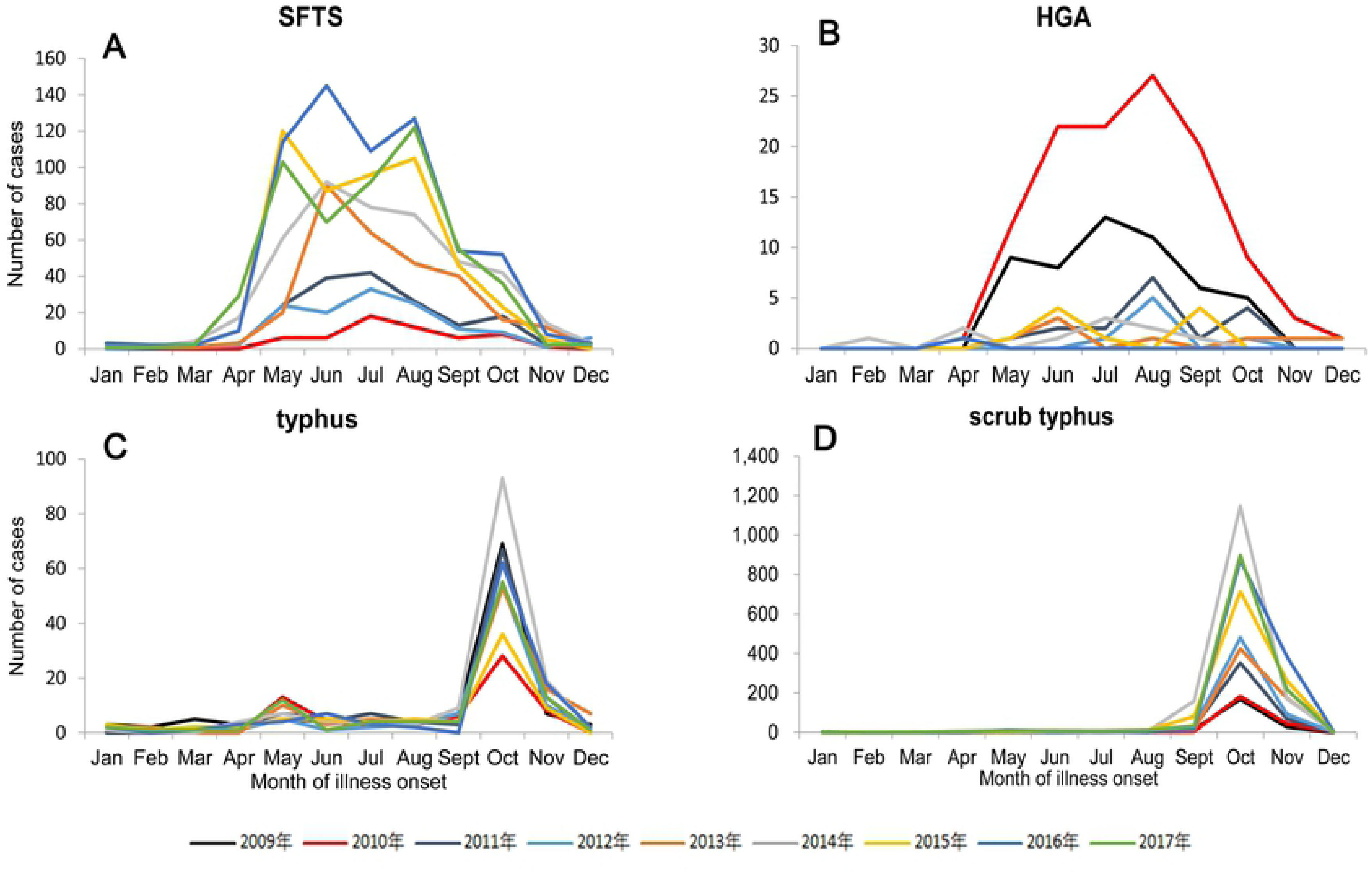
During 2009-2017, the aggregated number of cases by month in Shandong Province. (A): The aggregated number of SFTS cases by month from 2010 to 2017. (B): The aggregated number of HGA cases by month from 2009 to 2016. (C): The aggregated number of typhus cases by month from 2009 to 2017. (D): The aggregated number of scrub typhus cases by month from 2009 to 2017.

#### Population distribution

Of the total SFTS cases, 1384 cases were male and 1347 cases were female, the male-to-female ratio was 1.03: 1, there were slightly more male cases than female cases, but the difference was not statistically significant (χ^2^=0.012, P = 0.924) (Table 1). The majority of SFTS cases were farmers (86.01%, 2349/2731) and the occupation distribution of SFTS cases in different years were similar (Table 1; Fig 4A). 94.87% (2591/2731) of SFTS cases occurred in individuals aged over 40 years (Table 1). The highest peak of age group distribution of number of SFTS cases occurs in the 60-65 age group. However, the highest peak of the age group distribution of SFTS incidence rate lags behind 2 age groups and appears in the 70-75 age group (Fig 5A).

**Fig 4.**
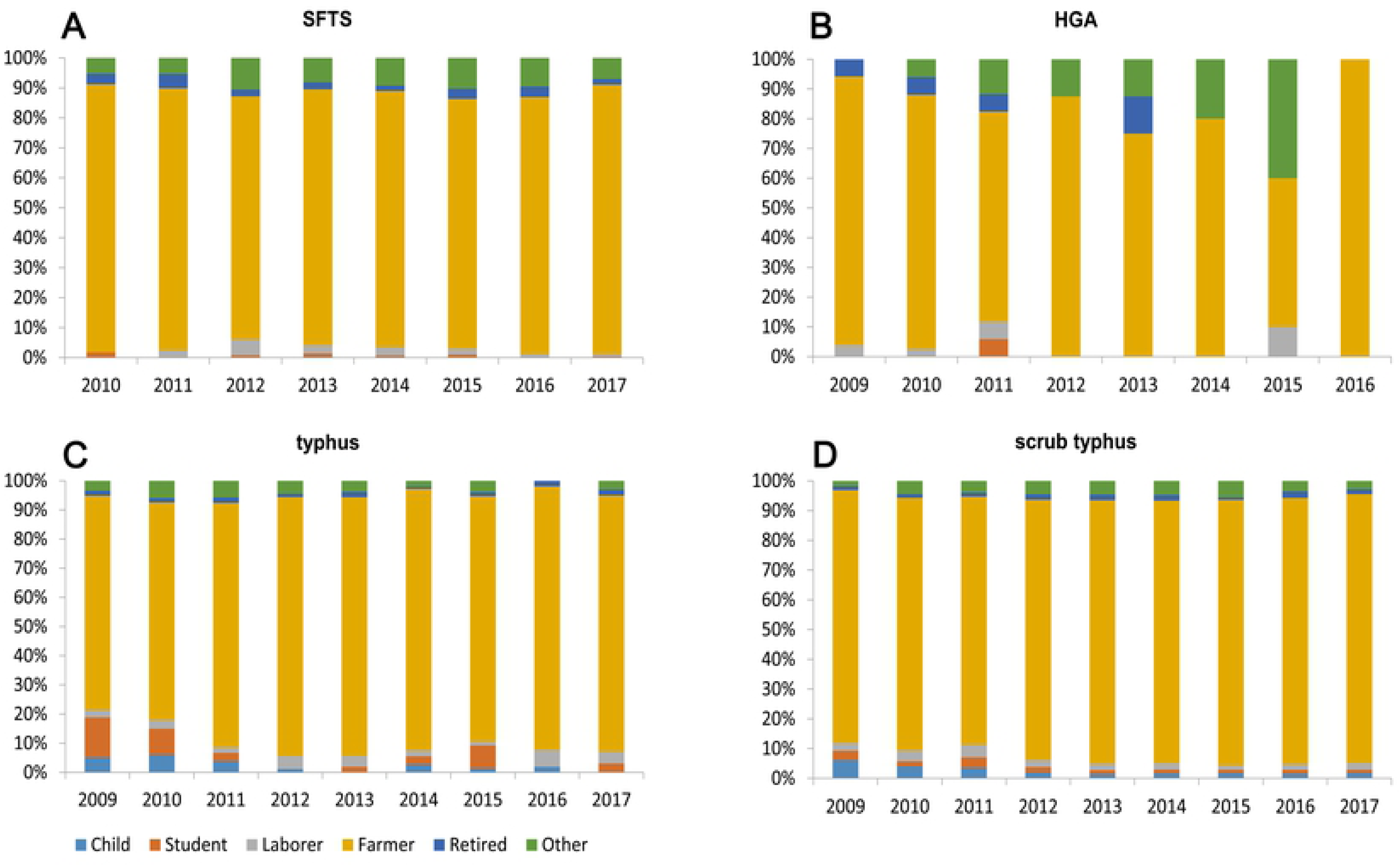
The proportion of different occupation of 4 species natural-focal diseases cases by year, 2009-2017. (A): Proportion of different occupations in SFTS cases by year in Shandong Province. (B): Proportion of different occupations in HGA cases by year in Shandong Province. (C): Proportion of different occupations in typhus cases by year in Shandong Province. (D): Proportion of different occupations in scrub typhus cases by year in Shandong Province.

**Fig 5.**
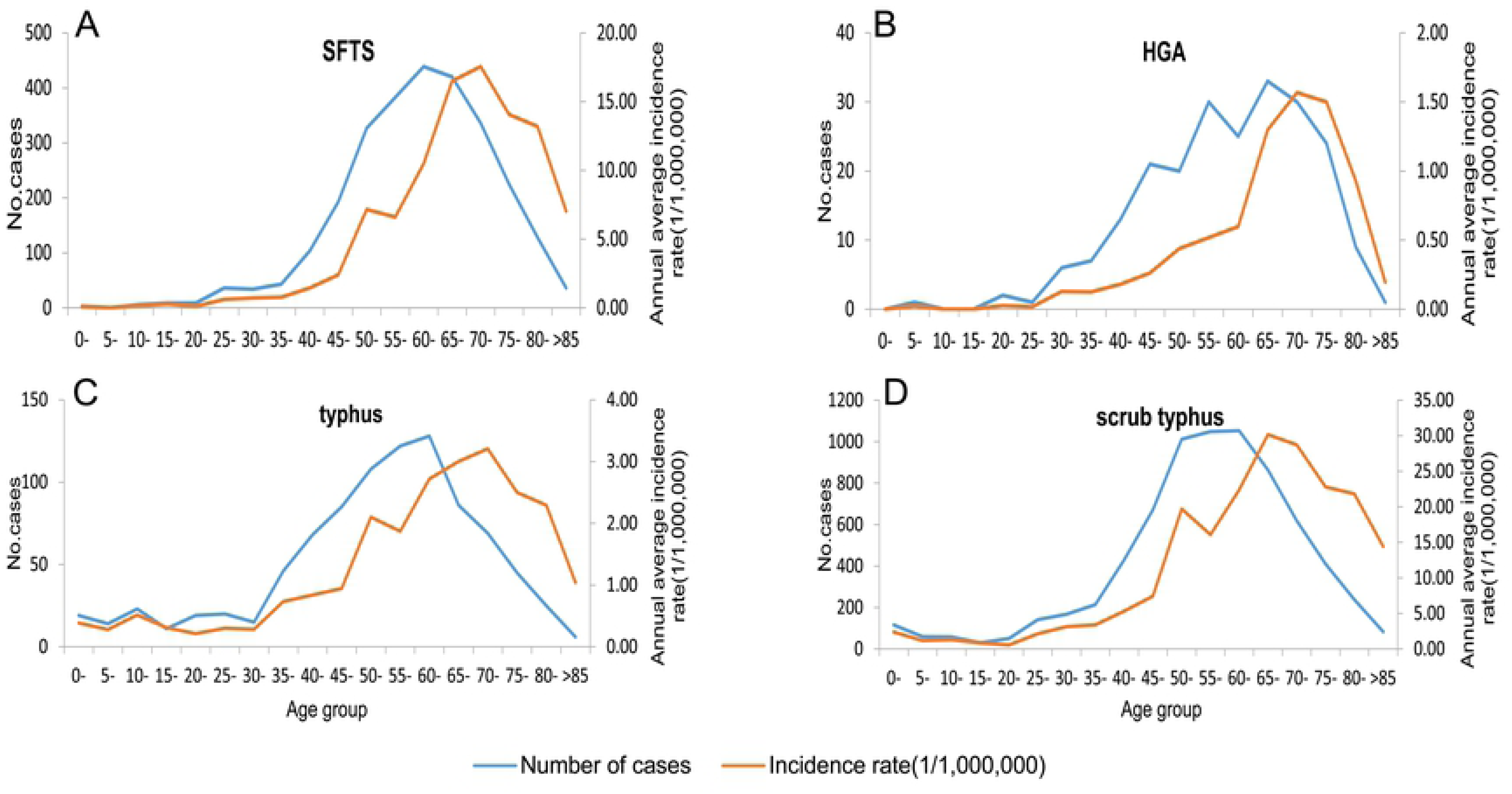
Age group distribution of the number of cases and incidence rate in Shandong, 2009-2017. (A): Age group distribution of SFTS cases and incidence rate. (B): Age group distribution of HGA cases and incidence rate. (C): Age group distribution of typhus cases and incidence rate. (D): Age group distribution of scrub typhus cases and incidence rate.

#### Regional distribution

During 2010–2017, 86.34% of SFTS cases were limited to 7 of 17 cities in Shandong Province: Yantai City (24.86%, 679/2731), Weihai City (16.33%, 446/2731), Tai’an City (12.45%, 340/2731), Jinan City (11.09%, 303/2731), Weifang City (8.75%, 239/2731), Linyi City (7.54%, 206/2731), Qingdao City (5.31%, 145/2731). Besides these 7 cities, other cities reported only 373 SFTS cases (Table 2). Of note, 46.50% (1270/2731) of SFTS cases were limited to Shandong Peninsula (including Yantai, Weihai and Qingdao). Furthermore, no case had been reported in Heze City during 2010 and 2017 (Table 2, Fig 6). The numbers of affected cities during 2010 and 2017 were 5, 11, 11, 15, 13, 14, 13 and 14, respectively (data not shown).

**Fig 6.**
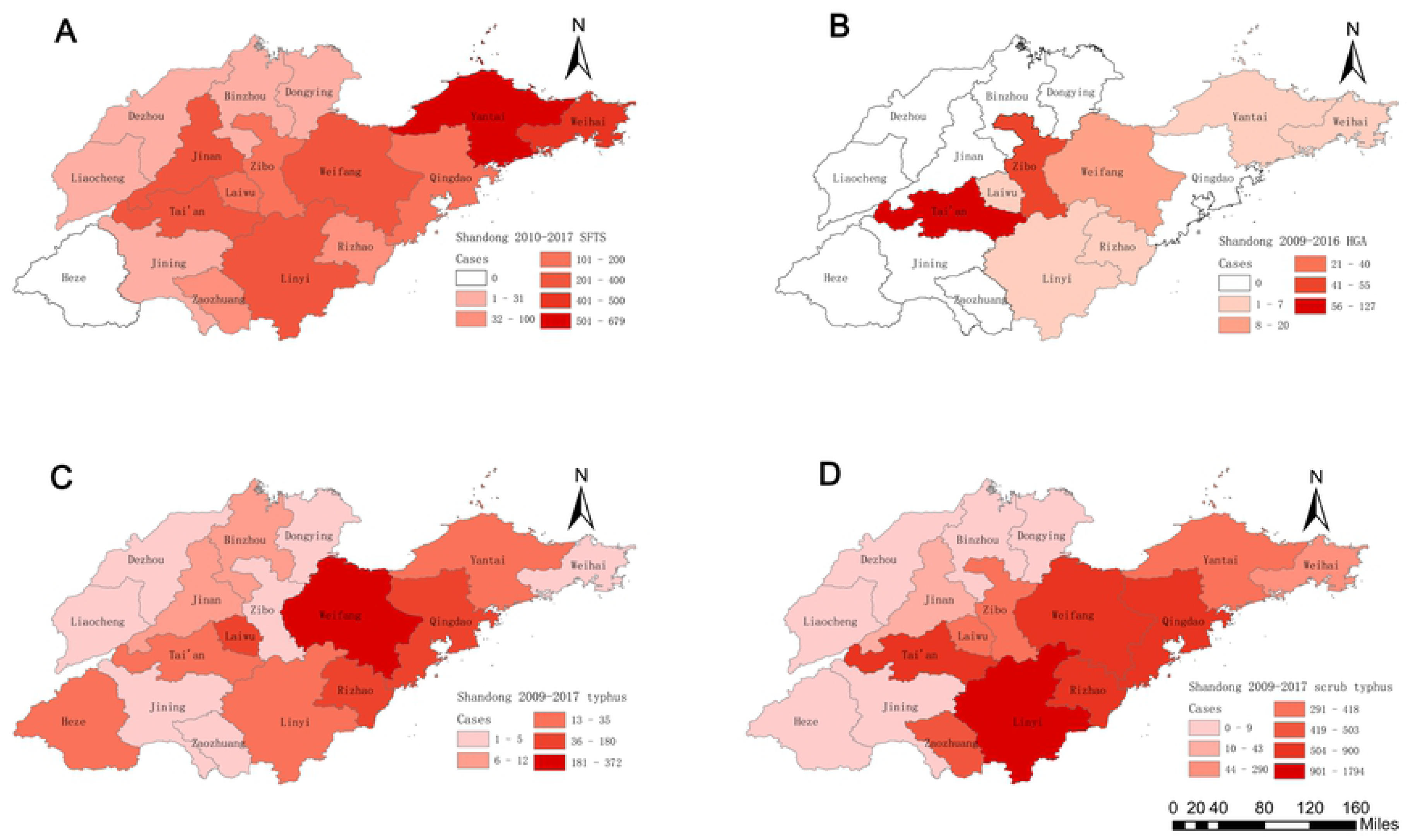
The geographic distribution of 4 species natural-focal diseases cases in Shandong, 2009-2017. (A): Geographical distribution of SFTS cases in Shandong Province. (B): Geographical distribution of HGA cases in Shandong Province. (C): Geographical distribution of typhus cases in Shandong Province. (D): Geographical distribution of scrub typhus cases in Shandong Province.

**Table 2.** Regional distribution of 4 species of natural-focal diseases in Shandong province, 2009-2017.

|  | SFTS<br>(2010-2017) |  | HGA<br>(2009-2016) |  | Typhus<br>(2009-2017) |  | Scrub typhus<br>(2009-2017) |  |
| --- | --- | --- | --- | --- | --- | --- | --- | --- |
|  | No. of<br>cases | Proporti<br>on (%) | No. of<br>cases | Proport<br>ion (%) | No. of<br>cases | Proportio<br>n (%) | No. of<br>cases | Proportio<br>n (%) |
| City |  |  |  |  |  |  |  |  |
| Qingdao | 145 | 5.31 | 0 | 0.00 | 99 | 10.89 | 835 | 11.50 |
| Yantai | 679 | 24.86 | 7 | 3.14 | 22 | 2.42 | 359 | 4.94 |
| Weifang | 239 | 8.75 | 20 | 8.97 | 372 | 40.92 | 815 | 11.23 |
| Weihai | 446 | 16.33 | 4 | 1.79 | 1 | 0.11 | 290 | 3.99 |
| Rizhao | 66 | 2.42 | 4 | 1.79 | 106 | 11.66 | 900 | 12.40 |
| Linyi | 206 | 7.54 | 2 | 0.90 | 30 | 3.30 | 1794 | 24.71 |
| Liaocheng | 1 | 0.04 | 0 | 0.00 | 1 | 0.11 | 1 | 0.01 |
| Laiwu | 117 | 4.28 | 4 | 1.79 | 180 | 19.80 | 418 | 5.76 |
| Dezhou | 11 | 0.40 | 0 | 0.00 | 1 | 0.11 | 0 | 0.00 |
| Dongying | 7 | 0.26 | 0 | 0.00 | 3 | 0.33 | 1 | 0.01 |
| Zibo | 118 | 4.32 | 55 | 24.66 | 1 | 0.11 | 390 | 5.37 |
| Binzhou | 8 | 0.29 | 0 | 0.00 | 12 | 1.32 | 8 | 0.11 |
| Zaozhuang | 38 | 1.39 | 0 | 0.00 | 5 | 0.55 | 503 | 6.93 |
| Jining | 7 | 0.26 | 0 | 0.00 | 2 | 0.22 | 9 | 0.12 |
| Heze | 0 | 0.00 | 0 | 0.00 | 35 | 3.85 | 1 | 0.01 |
| Jinan | 303 | 11.09 | 0 | 0.00 | 7 | 0.77 | 43 | 0.59 |
| Tai'an | 340 | 12.45 | 127 | 56.95 | 32 | 3.52 | 893 | 12.30 |

#### Death cases distribution

A total of 251 deaths were reported in Shandong Province from 2010 to 2017 and the fatality rate was 9.19%, with the highest recorded in 2011 (13.69%, 23/168) and the lowest in 2016 (7.00%, 44/629). The fatality rate declined with the year (Linear-by-linear Association, *χ*^2^=11.823, *P*=0.001). The fatality rate of male (10.04%, 139/1384) is slightly higher than female (8.61%, 116/1347), but the difference was not statistically significant (*χ*^2^=1.653, *P*=0.212). The fatality rates of the laborers, the retirees, the farmers, and the others are respectively 11.76% (6/51), 10.39% (8/77), 9.49% (223/2349), and 5.91% (14/237). All SFTS death cases occurred in individuals aged over 35 years, between 35 years and 85 years, and the fatality rate increased with age (Linear-by-linear Association, *χ*^2^=40.361, *P*<0.001), but the fatality rate declined in age groups of over 85 years old. In the 11 cities with SFTS deaths, there was a significant difference in the fatality rate (χ^2^=86.569, P<0.001). Dezhou had the highest fatality rate (18.18%, 2/11) and Weifang had the lowest (0.84%, 2/239), no deaths were reported in other 6 cities (Table 3).

**Table 3.** Demographic and epidemiological characteristics of fatality cases of 4 species natural-focal diseases in Shandong province, 2009-2017.

|  | SFTS<br>(2010-2017) |  | HGA<br>(2009-2016) |  | Typhus<br>(2009-2017) |  | Scrub typhus<br>(2009-2017) |  |
| --- | --- | --- | --- | --- | --- | --- | --- | --- |
|  | No. of<br>deaths | Fatality<br>rate (%) | No. of<br>deaths | Fatality<br>rate (%) | No. of<br>deaths | Fatality<br>rate (%) | No. of<br>deaths | Fatality<br>rate (%) |
| Age group (years) |  |  |  |  |  |  |  |  |
| 0- | 0 | 0.00 | 0 | 0.00 | 0 | 0.00 | 0 | 0.00 |
| 35- | 1 | 2.33 | 0 | 0.00 | 0 | 0.00 | 0 | 0.00 |
| 40- | 3 | 2.88 | 0 | 0.00 | 0 | 0.00 | 0 | 0.00 |
| 45- | 5 | 2.59 | 0 | 0.00 | 0 | 0.00 | 0 | 0.00 |
| 50- | 16 | 4.89 | 0 | 0.00 | 0 | 0.00 | 0 | 0.00 |
| 55- | 34 | 8.88 | 1 | 3.33 | 0 | 0.00 | 1 | 0.10 |
| 60- | 43 | 9.79 | 1 | 4.00 | 0 | 0.00 | 0 | 0.00 |
| 65- | 47 | 11.19 | 2 | 6.06 | 0 | 0.00 | 0 | 0.00 |
| 70- | 47 | 13.99 | 0 | 0.00 | 0 | 0.00 | 0 | 0.00 |
| 75- | 33 | 14.67 | 1 | 4.17 | 0 | 0.00 | 0 | 0.00 |
| 80- | 20 | 15.63 | 0 | 0.00 | 0 | 0.00 | 0 | 0.00 |
| >85 | 2 | 5.56 | 0 | 0.00 | 0 | 0.00 | 0 | 0.00 |
| Total | 251 | 9.19 | 5 | 2.24 | 0 | 0.00 | 1 | 0.01 |
| Gender |  |  |  |  |  |  |  |  |
| Male | 139 | 10.04 | 5 | 4.39 | 0 | 0.00 | 0 | 0.00 |
| Female | 116 | 8.61 | 0 | 0.00 | 0 | 0.00 | 1 | 0.02 |
| Occupation |  |  |  |  |  |  |  |  |
| Child | 0 | 0.00 | 0 | 0.00 | 0 | 0.00 | 0 | 0.00 |
| Student | 0 | 0.00 | 0 | 0.00 | 0 | 0.00 | 0 | 0.00 |
| Laborer | 6 | 11.76 | 0 | 0.00 | 0 | 0.00 | 0 | 0.00 |
| Farmer | 223 | 9.49 | 4 | 2.15 | 0 | 0.00 | 0 | 0.00 |
| Retiree | 8 | 10.39 | 1 | 8.33 | 0 | 0.00 | 0 | 0.00 |
| Other | 14 | 5.91 | 0 | 0.00 | 0 | 0.00 | 1 | 0.33 |
| Year |  |  |  |  |  |  |  |  |
| 2009 | - | - | 1 | 1.92 | 0 | 0.00 | 0 | 0.00 |
| 2010 | 6 | 10.53 | 2 | 1.71 | 0 | 0.00 | 0 | 0.00 |
| 2011 | 23 | 13.69 | 1 | 5.88 | 0 | 0.00 | 1 | 0.22 |
| 2012 | 17 | 12.78 | 1 | 12.50 | 0 | 0.00 | 0 | 0.00 |
| 2013 | 25 | 8.45 | 0 | 0.00 | 0 | 0.00 | 0 | 0.00 |
| 2014 | 58 | 13.33 | 0 | 0.00 | 0 | 0.00 | 0 | 0.00 |
| 2015 | 41 | 8.23 | 0 | 0.00 | 0 | 0.00 | 0 | 0.00 |
| 2016 | 44 | 7.00 | 0 | 0.00 | 0 | 0.00 | 0 | 0.00 |
| 2017 | 37 | 7.18 | - | - | 0 | 0.00 | 0 | 0.00 |
| City |  |  |  |  |  |  |  |  |
| Dezhou | 2 | 18.18 | 0 | 0.00 | 0 | 0.00 | 0 | 0.00 |
| Yantai | 99 | 14.58 | 0 | 0.00 | 0 | 0.00 | 0 | 0.00 |
| Jinan | 39 | 12.87 | 0 | 0.00 | 0 | 0.00 | 0 | 0.00 |
| Tai'an | 43 | 12.65 | 4 | 3.14 | 0 | 0.00 | 0 | 0.00 |
| Qingdao | 14 | 9.66 | 0 | 0.00 | 0 | 0.00 | 0 | 0.00 |
| Zibo | 9 | 7.63 | 0 | 0.00 | 0 | 0.00 | 0 | 0.00 |
| Weihai | 29 | 6.50 | 0 | 0.00 | 0 | 0.00 | 0 | 0.00 |
| Rizhao | 4 | 6.06 | 0 | 0.00 | 0 | 0.00 | 0 | 0.00 |
| Laiwu | 4 | 3.42 | 0 | 0.00 | 0 | 0.00 | 0 | 0.00 |
| Linyi | 6 | 2.91 | 0 | 0.00 | 0 | 0.00 | 1 | 0.06 |
| Weifang | 2 | 0.84 | 1 | 5.00 | 0 | 0.00 | 0 | 0.00 |
| Other | 0 | 0.00 | 0 | 0.00 | 0 | 0.00 | 0 | 0.00 |

**Note: “-” represents lack of data.**

### HGA

From 2009 to 2016, a total of 223 confirmed cases of HGA including 5 deaths in Shandong Province were reported to SDRIS.

#### Time distribution

There were 52 HGA cases in 2009 and 117 HGA cases in 2010 in Shandong Province, however, there were few HGA cases during 2011-2016 (Fig 2B). An average of 0.29 cases per one million residents each year were reported in Shandong Province, with the highest recorded in 2010 (1.22 cases/1,000,000) and the lowest in 2016 (0.01 cases/1,000,000) (Fig 2A). The incidence of HGA had obvious seasonal characteristics, with 94.62% (211/223) of the HGA cases clustering between May and October. The peaks of the reported HGA cases in 2009 - 2016 occurred in the summer season - July, August, August, August, June, July, June, respectively (Fig 3B; Table 1).

#### Population distribution

Of the total HGA cases, 114 cases were male and 109 cases were female, and the male-to-female ratio was 1.05: 1. There were slightly more male cases than female cases, but the difference was not statistically significant (χ^2^=0.027, P = 0.984) (Table 1). The majority of HGA cases were farmers (83.41%, 186/223) (Table 1; Fig 4B). 92.38% (206/223) of the HGA cases occurred in individuals aged over 40 years old (Table 1). The highest peak of age group distribution of number of HGA cases occurs in the 65-70 age group. However, The 65-70 age group had more HGA cases compared to the other age groups. The 70-75 age group showed the highest HGA incidence rate (Fig 5B).

#### Regional distribution

During 2009–2016, 90.58% of HGA cases were limited to 3 of 17 cities in Shandong Province: Tai’an City (56.95%, 127/223), Zibo City (24.66%, 55/223), Weifang City (8.97%, 20/223). Besides these 3 cities, 21 HGA cases were reported in other cities (Table 2; Fig 6). The numbers of affected cities during 2009 and 2016 were 5, 8, 2, 2, 3, 4, 1 and 1, respectively (data not shown).

#### Death cases distribution

A total of 5 deaths were reported in Shandong Province from 2009 to 2016 and the fatality rate was 2.24%. 1 case occurred in 2009, 2 cases occurred in 2010, and 1 case occurred in 2011, 1 case occurred in 2012. 5 cases of death were all male. The fatality rate of male (4.39%, 5/114) is higher than female (0.00%, 0/109), but the difference was not statistically significant (χ^2^=3.094, P=0.079). Among the 5 death cases, one was a retiree and the other 4 were farmers. All HGA death cases occurred in individuals aged over 55 years old. Only 2 cities had death cases, the fatality rate of Tai’an city was 3.14% (4/127), Weifang was 5% (1/20). No death was reported in 15 other cities (Table 3).

### Typhus

From 2009 to 2017, a total of 909 confirmed cases of typhus in Shandong Province were reported to SDRIS.

#### Time distribution

During 2007-2017, the number of cases of typhus in Shandong Province was relatively stable, with the highest recorded in 2014(146 cases) and lowest recorded in 2010(67 cases) (Fig 2B). An average of 1.04 cases per one million residents each year was reported in Shandong Province, with the highest recorded in 2014(1.49 cases/1,000,000) and the lowest in 2010 (0.70 cases/1,000,000) (Fig 2A). The incidence of typhus had obvious seasonal characteristics, 69.31% (630/909) of typhus cases occurred in October and November, and peaked in October each year (Fig 3C; Table 1).

#### Population distribution

Of the total typhus cases, 482 cases were male and 427 cases were female, and the male-to-female ratio was 1.13: 1. There were slightly more male cases, but the difference was not statistically significant (χ^2^=2.184, P = 0.144) (Table 1). The majority of typhus cases were farmers (85.04%, 773/909) (Table 1; Fig 4C). All age groups were susceptible to typhus, 81.63% (742/909) of the typhus cases occurred in individuals aged over 40 years old (Table 1). The highest peak of age group distribution of number of typhus cases occurs in the 60-65 age group. However, the highest peak of the age group distribution of typhus incidence rate lags behind 2 age groups and appears in the 70-75 age group (Fig 5C).

#### Regional distribution

During 2009–2017, 83.28% of typhus cases were limited to 4 of 17 cities in Shandong Province: Weifang City (40.92%, 372/909), Laiwu City (19.80%, 180/909), Rizhao City (11.66%, 106/909), Qingdao City (10.89%, 99/909). Besides these 4 cities, only 152 typhus cases were reported in other cities (Table 2; Fig 6). The numbers of affected cities during 2009 and 2017 were 10, 10, 8, 11, 10, 9, 13, 8 and 11, respectively (data not shown).

### Scrub Typhus

From 2010 to 2017, a total of 7260 confirmed cases of scrub typhus including 1 death in Shandong Province were reported to SDRIS.

#### Time distribution

During 2009-2014, the number of annual reported cases increased year by year, with the highest recorded in 2014 (1495 cases), from 2015 to 2017, the number of the annual cases was between 1093 and 1311 (Fig 2B). Of note, the number of cases increased massively by 866 from 2013 to 2014 with the annual growth rate of 137.68%. During 2009-2017, an average of 8.21 cases per one million residents were reported each year in Shandong Province, with the highest recorded in 2014 (15.27 cases/1,000,000) and the lowest in 2009 (2.40 cases/1,000,000) (Fig 2A). The incidence of scrub typhus had obvious seasonal characteristics, 97.06% (7047/7260) of scrub typhus cases were occurred between September and November, with the highest peak in October (Fig 3D; Table 1).

#### Population distribution

Of the total scrub typhus cases, 3148 cases were male and 4112 cases were female. The male-to-female ratio was 0.77: 1, and the difference was statistically significant (χ^2^=151.16, P<0.001) (Table 1). The majority of scrub typhus cases were farmers (88.43%, 6420/7260) (Table 1, Fig 4D). All age groups are susceptible, 88.53% (6427/7260) of typhus cases occurred in individuals aged over 40 years old (Table 1). The 60-65 age group had more scrub typhus cases compared to the other age groups but had a lower incidence rate than the 65-70 age group which showed the highest incidence rate among the groups (Fig 5D).

#### Regional distribution

During 2009–2017, the scrub typhus cases had been widespread in most of the cities in Shandong Province, 72.13% of scrub typhus cases were limited to 5 of 17 cities in Shandong Province: Linyi City (24.71%, 1794/7260), Rizhao City (12.40%, 900/7260), Tai’an City (12.30%, 893/7260), Qingdao City (11.50%, 835/7260), Weifang City (11.23%, 815/7260). 2023 scrub typhus cases were reported in other cities in Shandong Province (Table 2; Fig 6). The numbers of affected cities during 2009 and 2017 were 11, 12, 13, 12, 11, 13, 12, 13 and 13, respectively (data not shown).

#### Death cases distribution

A female from the 55-60 year old group in Linyi died from scrub typhus in 2011. This was the only death case reported in Shandong Province during 2009–2017 (Table 3).

## Discussion

In recent years, with the rapid advancement of urbanization in China and the change of people’s living style, the epidemic characteristics of natural-focal diseases had been also changing. In this study, surveillance data on 4 species natural-focal diseases was used to describe the epidemic characteristics of natural-focal diseases from 2009 to 2017 in Shandong, northern China.

### Annual distribution

The results showed that the number of cities with SFTS cases in Shandong Province increased rapidly from 5 in 2010 to 14 in 2017, and the number of SFTS has been on the rise in the mass except for 2012 and 2017 declined slightly from 2010 to 2017. Three factors may contribute to the result. First, since the first discovery of SFTSV in China in 2009, doctors and health departments were more and more awared of SFTS, and missed diagnosis or misdiagnosis was reduced[2]. Second, SFTSV might has been spread to more areas through humans, ticks, small mammals, or birds[7, 10]. It result in more people have opportunity to be infected with SFTSV. Third, with the advancement of China’s new rural construction and urbanization, the opportunities for ticks to contact people have increased. Comparison of the annual average incidence of Shandong and the annual average incidence from 2011 to 2016 of China, the incidence of Shandong (3.47 cases/1,000,000) is far higher than the national level(0.65 cases/1,000,000)[11]. The results show that Shandong is an important region for the epidemic area of SFTS in China. However, the reasons for this result need to be further studied.

Shandong as an emerging epidemic focus of HGA, the first confirmed case was found in 2008[12]. During 2009-2016, the number of cities with HGA cases has decreased from 8 in 2010 to 1 in 2016, the number of reported cases of HGA reached the highest in 2010, then sharply reduced. This may be related to the following factors. First, before 2010, many SFTS patients were misdiagnosed as HGA[13]. With the discovery of SFTSV and the improvement of SFTS diagnostic methods, the incidence of HGA tends to be true. Second, prevention and control measures of department of health were effective. Third, because there are not many cases in the past few years, the health department has relaxed its vigilance against HGA.

The incidence of typhus in Shandong Province has been very serious. From 1994 to 2003, 6653 cases of typhus occurred in Shandong Province, accounting for 14.10% (6653/47145) of the total number of cases in China[8]. However, the number of typhus cases in Shandong has decreased dramatically, 909 cases had occurred in 2009-2017 and none of them died, and the number of cities with typhus cases is relatively stable, about 10 per year in 2009-2017. This may be related to the improvement of people’s living environment and health habits in recent years, such as rural toilet improvement, garbage sorting and recycling, river pond treatment and other ecological environmental work in Shandong rural areas, which may reduce the chances of people direct contact with xenopsyllae cheopis and rodents. In addition, studies have shown that typhus can spread in international travel[14, 15]. With the increasing number of tourists abroad in China, monitoring of returned tourists should be strengthened. The number of cases of typhus in 2009-2017 is relatively stable, which may be related to the passive monitoring strategy of the Shandong Center for Disease Control and Prevention (Shandong CDC). According to our understanding, the passive monitoring strategy relies mainly on qualified units (hospitals and CDCs) to conduct direct network reporting. Due to the low fatality rate of the disease, it has been neglected by many units, and the units that have been reported on the initiative are relatively few and fixed, so the number of cases of typhus has been relatively stable.

Shandong Province as an emerging epidemic focus of scrub typhus, the number of reported cases in where has increased rapidly since the first cases reported in Shandong Province in 1986[16-19]. This study shows that the number of cities with scrub typhus cases is relatively stable and the number of annual reported cases of scrub typhus increased slowly before 2013, but the sharp increase from 2013 to 2014 and then has been maintained at a high level. This may be related to the following factors. First, it is understood that Shandong CDC conducted active surveillance of scrub typhus in parts of five cities (Linyi, Zibo, Tai’an, Qingdao, Laiwu) from April 2013 to December 2015, which may cause a sharp increase in the number of scrub typhus cases after 2013. Second, with the advancement of China’s new rural construction and urbanization, the opportunities for chigger mites to contact people have increased. Third, distribution range of vector chigger mites and main reservoir host rodents, availability of detection facilities as the increasing investment of health resources all may affect the number of annual reported cases.

### Monthly distribution

The incidence of 4 species natural-focal diseases all has obvious seasonal. May to August is the epidemic peak of SFTS and HGA in Shandong Province, which was similar to other SFTS epidemic areas, such as those in Henan, Anhui, Jiangsu, and Zhejiang provinces[20-22]. Haemaphysalis longicornis is the main tick species in Shandong Province, SFTSV and AP are believed to be mainly transmitted by Haemaphysalis longicornis ticks bite in Shandong Province[7, 23]. Due to the lack of data on tick density fluctuation in Shandong Province, we refer to two adjacent provinces, Henan and Jiangsu, which are similar to the geographical location, climate type and tick species distribution of Shandong Province. The peak of tick density in Henan and Jiangsu occurs from May to August, and we speculate that the peak of tick density in Shandong Province also occurs from May to August[24, 25]. Therefore, we believe that the epidemic peaks of SFTS and HGA are consistent with the fluctuation of tick density in Shandong Province.

The epidemic peak of typhus in Shandong was from October to November, which was different from other typhus epidemic areas, such as those in Henan, Yunnan, and Zhejiang provinces[26-28]. The reason for this may be that Shandong is located in the north of these epidemic areas, with lower temperature, result in the main vector of Xenopsylla cheopis to reach the number of peak later. In addition, October and November were the harvest time in Shandong Province, and farmers had frequent contacts with the Xenopsylla cheopis may resulted in an increase in the number of cases.

September to November is the epidemic peak of scrub typhus, which may be associated with the following factors. First, September to November is the harvest time in Shandong Province, increasing outdoor activities of farmers during this period and enhanced the risk for farmers to contact with the mainly transmission vector of Leptotrombidium scutellare[29]. Second, rodents and Leptotrombidium scutellare as the main reservoir host and transmission vector for Orientia tsutsugamushi in Shandong Province. Their density fluctuations are closely related to the seasonal distribution of scrub typhus cases, and the monthly distribution of scrub typhus cases was consistent with the fluctuation of Leptotrombidium scutellare[9, 30, 31]. Third, some studies have shown that meteorological factors affect the incidence of scrub typhus, such as temperature, sunlight, precipitation, etc[16, 32, 33]. The climatic conditions of Shandong Province from September to November may be most suitable for the occurrence of scrub typhus. The above factors may contribute to the high infection rate of scrub typhus during autumn.

### Population distribution

The high-risk groups of the 4 species diseases all were farmers and the elderly. This may be related to the following factors. First, with the advancement of China’s urbanization process and the expansion of enrollment in higher education institutions, a large number of middle-aged men and young people from rural areas have entered the city to work or study, and the elderly become the main force of agricultural production[34]. Therefore, elder people who had more opportunity to expose to insect vector when undertaking farm work than young people who work or study in cities except Spring Festival[35]. Second, the immune function of the elderly may be lower than young people. If they are infected with these 4 species diseases, elderly may get severe diseases and go to hospital for treatment[36, 37]. It is thereby that more elderly cases were identified.

The highest peaks of the age group distribution of the incidence rate of the four diseases lag behind the highest peak of the age group distribution of the number of cases 1 or 2 age groups. The 60-65 age group had the highest number of cases (except HGA), but the 70-75 age group had the highest incidence rate (except scrub typhus). That is related to the age structure of the population in Shandong Province. Compared with the 60-65 age group, the 70-75 age group has a smaller population base, so the incidence rate is higher. Therefore, this may also reflect that the 70-75 age group is the highest risk group for natural-focal diseases in Shandong Province.

The results showed that the incidence of scrub typhus among females was higher than that observed in males, which is consistent with previous studies[17, 18]. Two factors may contribute the result. First, the increased proportion of females engaging in outdoor activities[38]. Second, female may be more susceptible to Orientia tsutsugamushi than males[39]. This requires further experimentation to explore.

### Regional distribution

SFTS cases in Shandong are mainly concentrated in Mount Yimeng (Linyi, Jinan, Tai’an, Weifang, Zibo, Laiwu) and Shandong peninsula (Qingdao, Yantai, Weihai). HGA cases in Shandong are mainly concentrated in Mount Yimeng (Tai’an, Weifang, Zibo). Mount Yimeng is located in the middle-southern part of Shandong Province, Laoshan is located in the Shandong Peninsula. The density of ticks may be higher in the mountains area, where shrub grassland is abundant, so there were more SFTS and HGA cases[40]. Therefore, the mountainous areas with abundant vegetation cover should be the key areas for SFTS and HGA prevention and control. In addition, it should be noted that SFTS and HGA cases are not reported in some areas does not mean that the area must not have occurred. It may be because the patient’s clinical performance is not serious and there is no medical diagnosis or doctor misdiagnosis.

Typhus cases in Shandong are mainly concentrated in mountainous inland (Laiwu) and coastal areas (Rizhao, Qingdao, Weifang). The incidence of typhus cases may be related to rural rodent density and Xenopsylla cheopis density in these areas, but the relevant survey data are lacking at present. Subsequent studies can further explore the factors affecting the distribution of typhus cases in different regions by investigating the density of rural rodent density and Xenopsylla cheopis density in different regions.

Most of scrub typhus report cases in Shandong Province were concentrated in mountainous inland (Linyi, Tai’an) and coastal hilly areas (Rizhao, Qingdao, Weifang), which can be partly explained by the chigger mites population abundance[41]. It is possible that chigger mites are more abundant in the low mountains and hills because of the rich vegetation cover in the low mountains and hills, the overgrown weeds in the mountains, the suitable temperature, the abundant precipitation and the humid environment, it is suitable for the survival and reproduction of chigger mites.

### Death cases distribution

All SFTS death cases occurred in individuals aged over 35 years, and laborer, retiree, and farmer have a higher fatality rate. Two factors may contribute these results. First, at this age groups, most of the cases are farmers and laborers, and their medical conditions are poor in rural areas or the construction site where they live lead to the diagnosis and treatment may be delayed. Combined studies show that the time spent in diagnosis affects prognosis[42]. Second, retirees are elderly, if infected by SFTSV, elderly may get severe diseases[36]. So the fatality rate is higher than that of mild or even asymptomatic patients.

Previously studies have reported that fatality rate of SFTS increases with age[11, 43]. Our study also indicated that age was closely related to the SFTS fatality rate. The decrease in fatality in the age group over 85 may be due to the fact that the number of cases is too small and unrepresentative, which is a contingency event. Some factors associated with age including decreased immune function and comorbidities with chronic diseases may be relative to SFTS fatal outcome.

The fatality rate of SFTS had decreased from 2010 to 2017, but it remained very high in 2017(7.18%). The high fatality rate suggest that it is urgent to study effective vaccines and treatments for SFTS. The case fatality rate declined may be explained by doctors’ improvement in the level of diagnosis and treatment of SFTS cases.

The fatality rate of SFTS varies in different regions. This may be related to the following factors. First, the proportion of severe cases in different regions is different, but the lack of relevant data can’t be judged. Second, there are differences in medical level and doctor’s experience. Based on what we know, we believe that the latter is more likely to be the main reason.

All HGA death cases occurred in individuals aged over 55 years. Two factors may contribute the result. First, at this age groups, most of the cases are farmers, and their medical conditions are poor in rural areas where they live lead to the diagnosis and treatment may be delayed[44]. Second, it may be related to the low immune function of the elderly lead to the therapeutic effect is worse[37, 44]. The average case fatality rate was 2.24% in Shandong Province, which is higher than 0.3% in the United States[45]. It may be related to the two fact. First, at present, HGA still not recognized by many Chinese doctors, which is likely to cause misdiagnosis and delayed antibiotic therapy[46, 47]. Second, the outer membrane protein msp2/p44 as important virulence factors of AP pathogens, and their molecular characteristics are significantly different in strains isolated from China and the United States at the nucleic acid, amino acid and protein levels [48]. Therefore, Chinese patients often show more serious clinical manifestations and a higher mortality rate[48].

The five deaths of HGA were all concentrated in 2009-2012, which may be related to the fact that many SFTS were misdiagnosed as HGA during this time, which led to the emergence of death cases[13, 46].

Although only a scrub typhus death case occurred in Shandong Province from 2009 to 2017, this suggests that we were still unable to relax our vigilance in the treatment of patients with scrub typhus.

### Limitations

There are some limitations in our study. First, since the case data were obtained from a passive surveillance system, the reporting system might miss some cases because some patients have no clinical manifestations or have mild performance and have not been diagnosed and treated in hospital. In addition, many grassroots areas do not have the equipment and technology to diagnose and identify pathogens of natural-focal diseases. These factors may cause the number of reported cases to be much smaller than the actual number of cases. Second, due to the limitations of the data, the 4 species natural-focal diseases described are not wholly representative of natural-focal diseases in Shandong. Third, the residential address information of the reported cases is only accurate to the city-level, and it would be more precise to describe the regional distribution characteristics of the cases if it was accurate to the county and even town levels. This is more conducive to the accurate allocation of resources.

## Conclusions

Despite the limitations stated above, our study described the epidemic characteristics, and identified spatiotemporal clusters of 4 species natural-focal diseases (SFTS, typhus, scrub typhus and HGA) in Shandong Province during 2009–2017. Our findings may can contribute to public health officials allocate resources reasonably for natural-focal diseases prevention and control in Shandong Province.

## Supporting Information

**S1 Checklist: STROBE Checklist**

**S2 Appendix. Diagnosis criteria of 4 species natural-focal diseases.**

## Acknowledgements

We thanks staff members at the hospitals, local health departments, county-, district-, prefecture-, and provincial-level CDCs for assistance in data collection.

